# Similarity in Greater Chinese ideology is associated with intersubject neural similarity during viewing of a pro-China video

**DOI:** 10.64898/2026.07.25.740672

**Authors:** Po-Yuan Alan Hsiao, Feng-Chun Ben Chou, Pin-Hao Andy Chen

## Abstract

When people watch the same political narrative, they often form different interpretations. Although prior research has shown that political ideology can shape shared neural responses to sociopolitical content, less is known about how individual differences in national identity ideology are associated with inter-individual similarity in neural dynamics during naturalistic viewing, particularly in sociopolitical contexts where national identity is highly salient in public discourse. We examined Greater Chinese ideology, one dimension of national identity ideology, in relation to positive engagement and neural dynamics during viewing of the same pro-China video across two samples in Taiwan. In Study 1 (N = 52), higher Greater Chinese ideology was associated with greater positive engagement. In Study 2 (N = 60), participants underwent fMRI while viewing the same video. Intersubject representational similarity analysis showed that greater similarity in participants’ Greater Chinese ideology item-response profiles was associated with greater similarity in neural dynamics in regions associated with the frontoparietal control, salience, and default mode networks. Notably, the within-versus-between neural similarity effect was stronger among participants with relatively lower Greater Chinese ideology scores than among those with relatively higher scores. These findings link Greater Chinese ideology to positive engagement and similarity in neural dynamics during viewing of this pro-China video. Further research is needed to determine whether similar associations can be observed across other dimensions of national identity ideology and different political narratives.

## Introduction

Ideologies refer to the belief systems people use to make sense of society and to guide how they relate to others, and they vary systematically across individuals in ways that shape interpretation and evaluation of social information (Jost, 2017; Jost et al., 2008; Knight, 2006; Zmigrod, 2022). Understanding ideology is important because ideology influences how individuals perceive social reality, evaluate political information, and respond emotionally to group-related narratives (Jost et al., 2008; Waldfogel et al., 2021). These differences in interpretation and evaluation suggest that ideological variations may also be reflected in differences in neural responses to sociopolitical content (van Baar & FeldmanHall, 2022). One especially consequential form is national identity ideology, as it shapes how individuals define group belonging, interpret shared history, and position their community within a broader geopolitical landscape (L.-L. Huang, 2007; Smith, 1993; Zmigrod et al., 2018). Although past research has made substantial progress in characterizing the neural correlates of political ideology (de Bruin et al., 2023; Leong et al., 2020; van Baar & FeldmanHall, 2022; Zmigrod & Tsakiris, 2021), national identity ideology has received comparatively limited attention in neuroimaging research. Importantly, national identity ideology is not categorical but inherently graded, and individuals can occupy different positions along this continuum in ways that systematically shape how the same sociopolitical information is interpreted and evaluated (Mader & Schoen, 2023; Meeus et al., 2010). If national identity functions as a cognitive and affective lens, its influence should therefore be expressed not only in average neural responses, but also in variations across individuals exposed to the same ideology-related stimuli.

A key open question, then, is whether ideology-related neural similarity extends to national identity ideology. Most previous research has focused on partisan affiliation or liberal-conservative political ideology (Leong et al., 2020; van Baar et al., 2021), leaving the role of national identity less understood. National identity reflects how individuals understand their sense of national belonging and their relationship to a broader national community. Taiwan provides an informative context for examining this question because national identity varies considerably across individuals living within the same sociopolitical environment. These differences have been shaped by Taiwan’s historical and political development, including postcolonial legacies (F.-C. I. Yang & Mak, 2021), the societal memory of the martial law era (Kunze, 2024), democratization-related changes (W.-L. Chen et al., 2023), and persistent cross-strait tension (Chu, 2004; Jacobs, 2013; Wu, 2005). One important dimension of these differences is Greater Chinese ideology, which emphasizes shared history and cultural roots with mainland China and often views Taiwan as part of a broader Chinese nation (L.-L. Huang et al., 2004; L.-L. Huang, 2007). Individual differences in Greater Chinese ideology may, in turn, be associated with how individuals interpret and evaluate a pro-China political narrative. Individuals with different levels of Greater Chinese ideology may therefore experience the same narrative differently, particularly in how much they like and resonate with its content. This allows us to examine whether national identity ideology is associated with subjective engagement when individuals encounter the same political narrative.

Building on this question, a growing body of neuroimaging work has suggested that ideology can shape how people perceive, evaluate, and interpret sociopolitical information as it unfolds (Panish & Nam, 2023; Zheng & Han, 2025; Zmigrod, 2021; Zmigrod et al., 2021). When people encounter the same speeches or media narratives, ideological commitments can be associated with differences in attention and interpretation, as well as differences in affective engagement (Leong et al., 2020; Mills et al., 2015; Shteynberg et al., 2016; van Baar et al., 2021). Such differences have been linked to activity in affective and valuation systems including the amygdala, insula, and ventromedial pre-frontal cortex (Kaplan et al., 2016; Schreiber et al., 2013; Westen et al., 2006). Ideology may also shape what people do with these perceptions and feelings. Individuals often arrive at different judgments and decisions when exposed to the same sociopolitical information, consistent with differential recruitment of systems involved in cognitive control, belief updating, and conflict monitoring (Friedman & Robbins, 2022; Huber et al., 2015; Schulreich & Schwabe, 2021). Understanding these mechanisms matters because they relate to real-world outcomes, including political preferences and decision-making (Fournier et al., 2020; Jenke & Huettel, 2016; Jost et al., 2022). Together, these works support a general view that ideology can operate as an interpretive lens, shaping real-time meaning construction and affective evaluation during narrative exposure.

Naturalistic neuroimaging work further suggests that similarity in ideology can be expressed as shared neural dynamics when individuals attend to the same sociopolitical narratives (van Baar & FeldmanHall, 2022). When the content is ideologically diagnostic, individuals with similar identities or attitudes often show more similar neural dynamics in the brain networks linked to social cognition and cognitive control, such as the default mode network and the frontoparietal control network (Broom et al., 2022; Leong et al., 2020; van Baar et al., 2021). In naturalistic settings, these networks may support how viewers integrate unfolding information into higher-level representations of the narrative, with individual differences in experience and background contributing to variation in neural dynamics across individuals (L. J. Chang et al., 2021; de Bruin & FeldmanHall, 2025; Hakonen et al., 2022; Sievers et al., 2024; Yeshurun et al., 2021). Consistent with this view, Hakonen et al. (2022) also found that individuals who shared a cultural family background showed greater intersubject neural similarity while listening to the same narrative. However, most previous studies of ideology-related neural similarity have focused on partisan affiliation or liberal-conservative political ideology. It therefore remains unclear whether this relationship extends to national identity ideology, where individual differences concern how people understand national belonging rather than partisan or policy divisions. Taiwan provides an informative context for examining this question because national identity varies meaningfully among individuals living within the same sociopolitical environment (Lyu & Zhou, 2023; Zhong, 2016). We did not expect a qualitatively different relationship between ideological similarity and neural similarity in this context. Rather, we expected greater similarity in Greater Chinese ideology to be associated with greater similarity in neural dynamics while viewing the same pro-China narrative. This context provides a clear setting for investigation of whether ideological variation shapes neural responses in brain networks involved in narrative interpretation and evaluation, such as the default mode network and the frontoparietal control network, rather than in early sensory areas that primarily track stimulus-driven audiovisual features.

Our study addresses this gap by examining how individual differences in Greater Chinese ideology are associated with differences in neural dynamics during naturalistic viewing. We conducted two independent studies using the same set of videos from a recently published work (Chou et al., 2025), while focusing our hypotheses on examining the behavioral ratings and neural responses to a pro-China video that depicts China’s national development. Study 1 provided a behavioral test of whether individual differences in Greater Chinese ideology were associated with subjective engagement experience during naturalistic viewing of the pro-China video. To examine the individual differences in engagement experience, participants rated how much they liked the video and how much they felt it resonated using a visual analog scale, which were combined to generate a positive engagement score of the video. We hypothesized that individuals with higher Greater Chinese ideology would show higher positive engagement scores for the pro-China video. Study 2 extended the above study to the neural level using an independent sample scanned with fMRI while viewing the same set of videos. We tested whether similarity in Greater Chinese ideology item-response profiles was associated with similarity in neural dynamics during viewing of the pro-China video. To do so, we applied intersubject representational similarity analysis (ISRSA) (P.-H. A. Chen et al., 2020; Chou & Chen, 2025), which utilizes second order statistics similar to RSA (Kriegeskorte et al., 2008; Kriegeskorte & Douglas, 2018) to investigate whether intersubject similarity in national identity ideology item-response profiles was associated with intersubject similarity in neural dynamics during naturalistic viewing (P.H. A. Chen & Qu, 2021; Finn et al., 2020; Hsiao et al., 2024; Nguyen et al., 2019). We hypothesized a positive association between ideology item-response profile similarity and neural dynamic similarity, with the strongest associations expected to be found in the brain regions within the frontoparietal control network and the default mode network. Although guided by the theoretical expectations outlined above, the present studies used correlational analyses of available behavioral and neuroimaging data to characterize the associations of interest. Neither study was preregistered, and the hypotheses were not preregistered in advance. Although the analyses were guided by the theoretical expectations outlined above, they should therefore be considered theoretically informed but exploratory.

## Method

### A. Study 1

#### A.1. Participants

Study 1 used the independent video-rating dataset previously described by Chou et al. (2025). All 52 participants in the present study were included in the independent rating sample reported in that article. Chou et al. (2025) analyzed pandemic- and death-related ratings from this dataset for stimulus validation, whereas the present study examined the previously collected Greater Chinese ideology scores and liking and resonance ratings. The ideology–engagement analyses reported here were not included in the previous publication. Participants were recruited through online advertisements targeting university students who had not previously participated in the experiment. Data were collected between May 1st and May 21st, 2022. Sixty questionnaire records were received; after incomplete and duplicate records were removed, 52 participants who completed the study were retained. All participants were Taiwanese and resided in Taiwan. Most were university students at National Taiwan University, and Mandarin was the first language of most participants. Participants self-reported their gender using three response options: male, female, and non-binary. The final sample included 22 males and 30 females, with a mean age of 20.15 years (*SD* = 1.23, range = 18–24). None of the participants had a history of psychiatric or neurological disorders. All participants received monetary compensation for their participation and provided informed consent following the guidelines established by the Research Ethics Committee at National Taiwan University under approval number 202206HM013.

#### A.2. Experimental procedure and Stimuli

Participants watched a fixed sequence of fifteen videos that covered a broad range of everyday topics, including national ideology, nature, food, and music (Chou et al., 2025). Videos were separated by a 20-second fixation period to reduce carryover effects from the preceding clip. After each video, participants rated their subjective experience on six visual analog scales (0 = not at all, 100 = very much), assessing how much they liked the video, how resonant it felt, how much they would be willing to share it, the extent of positive emotion, the extent of negative emotion, and how well they understood the content. In the present study, we focused our analyses on the liking and resonance ratings, as these measures most directly captured positive engagement with each video clip.

The pro-China video was a Mandarin-language political video titled *Today’s China, As You Wish*, produced jointly by the Publicity Department of the Central Committee of the Communist Youth League of China and the Beijing Municipal Committee of the Communist Youth League. Released on January 8, 2016, the video commemorated the 40^th^ anniversary of Premier Zhou Enlai’ s death. According to contemporary reporting by *China Youth Daily*, the video was produced as part of the Communist Youth League’ s efforts to promote ideological convictions among young people through new media. The video is publicly available online (https://www.youtube.com/watch?v=ZMteHL9AkQY). To assess stimulus specificity, we also included a neutral control clip. The control video was a nature clip lasting 1 minute and 31 seconds depicting natural landscapes without narration or ideological messaging. This contrast allowed us to test whether associations between ideology and positive engagement experience were specific to the ideologically charged narrative rather than reflecting a general tendency to engage with audiovisual content. In addition, for the neuroimaging analyses in Study 2, we included a second non-political control video, Farewell, a 4 minute and 31 second short film depicting a father guiding his son to say goodbye to his deceased mother. Unlike the nature clip, Farewell contained an unfolding social narrative but no political or national-identity content. This contrast allowed us to test whether associations between ideology and positive engagement experience were specific to the ideologically charged narrative rather than reflecting a general tendency to engage with audiovisual content.

#### A.3. National identity ideology scale

After completing the video ratings, participants completed the national identity ideology scale (L.-L. Huang, 2007). This scale includes four subscales, namely Greater Chinese ideology, Kuomintang (KMT) legitimacy, Separation ideology, and Taiwanese refinement ideology. The scale comprised 34 items rated on a six-point agreement scale (1 = strongly disagree, 6 = strongly agree) and included four subscales: Greater Chinese ideology (11 items), KMT legitimacy (7 items), Separation ideology (11 items), and Taiwanese refinement ideology (5 items). The primary construct of interest was Greater Chinese ideology, which reflects endorsement of a shared cultural heritage and historical continuity with mainland China and often frames Taiwan as part of a broader Chinese nation (L.-L. Huang et al., 2004; L.-L. Huang, 2007). As a control construct of ideology, we also examined Taiwanese refinement ideology, which emphasizes a distinct Taiwanese identity with political and cultural autonomy (L.-L. Huang et al., 2004; L.-L. Huang, 2007). Including this control subscale allowed us to assess whether any association with positive engagement experience was specific to the theoretically focal ideology dimension. Although conceptually related, Greater Chinese ideology and Taiwanese refinement ideology were treated as distinct dimensions of national identity ideology rather than opposite endpoints of a single continuum. In contrast, KMT legitimacy primarily concerns the historical and political legitimacy of KMT rule, whereas Separation ideology concerns Taiwan’s political and cultural differentiation from China. These dimensions were not part of our focal hypotheses and were therefore not included in the primary analyses.

Internal consistency for Greater Chinese ideology, KMT legitimacy, Separation ideology, and Taiwanese refinement ideology was *α* = .86, .74, .75, and .73, respectively, in Study 1, and *α* = .88, .86, .82, and .69, respectively, in Study 2. The corresponding means (SDs) were 3.03 (0.72), 2.59 (0.73), 4.08 (0.58), and 4.64 (0.76) in Study 1, and 2.72 (0.83), 2.38 (0.93), 4.07 (0.69), and 4.56 (0.66) in Study 2. Complete item wording and subscale membership are provided in Table S1, with additional descriptive statistics in Table S2 and score distributions in Figure S1 in the Supplementary materials. Complete source-language item wording and subscale membership are reported in Table S1; additional descriptive statistics are reported in Table S2, and score distributions are shown in Figure S1.

#### A.4. Testing the association between national identity ideology and positive engagement experience

For each participant, we computed a Greater Chinese ideology score and a Taiwanese refinement ideology score by averaging items within each subscale. To quantify positive engagement with the pro-China video, we computed an engagement score as the mean of the liking and resonance ratings for the pro-China video. Positive engagement was defined as the average of liking and resonance ratings. These ratings were strongly correlated in both studies (Study 1: *r* = .81, 95% CI [.69, .89], *p* < .001; Study 2: *r* = .85, 95% CI [.76, .91], *p* < .001), supporting their combination into a single measure of positive engagement. To directly compare the associations of Greater Chinese ideology and Taiwanese refinement ideology with positive engagement during the pro-China video, we used Williams’ test for dependent correlations. We then conducted the same correlation analyses for the nature control video and used Williams’ test to compare the association between Greater Chinese ideology and positive engagement across the pro-China and control videos.

### B. Study 2

#### B.1. Participants and experimental procedure

Study 2 recruited an independent sample through online advertisements, targeting participants who had not previously participated in Study 1. Data were collected between November 12, 2021, and May 7, 2022. All participants underwent MRI safety screening before scanning. A total of 60 participants were scanned, and all participants were included in the final analysis. No participants were excluded on the basis of head motion or fMRI data quality. All participants were Taiwanese and resided in Taiwan. Most were university students at National Taiwan University, and Mandarin was the first language of most participants. Participants self-reported their gender using three response options: male, female, and non-binary. The final sample included 29 males and 31 females, with a mean age of 21.95 years (*SD* = 2.05, range = 18– 27). Fifty-eight participants were right-handed and two were left-handed. No participants reported a history of psychiatric or neurological disorders. Study 2 included two phases. Participants first underwent fMRI while viewing the same set of fifteen videos in the scanner. Participants then completed post-viewing ratings and the same national identity ideology scale outside the scanner. The video stimuli and national identity ideology scale were the same as in Study 1, but the post-viewing ratings differed slightly between the two studies. Both studies assessed liking, resonance, willingness to share, and comprehension. Study 1 included separate ratings of positive and negative emotion, whereas Study 2 used a single rating of overall emotional valence. All participants received monetary compensation for their participation and provided informed consent following the guidelines established by the Research Ethics Committee at National Taiwan University under approval number 202206HM013.

#### B.2. Image acquisition and preprocessing

##### Imaging parameters

MRI scanning was conducted using a 3T Siemens Prisma scanner equipped with a 32-channel head coil at the Imaging center for Integrated Body, Mind and Culture Research located at National Taiwan University. Structural images were acquired through the T1-weighted MPRAGE protocol (TR=2300ms, TE=2.32ms, voxel size=0.9×0.9×0.9mm, flip angle=8 degrees). Functional images were obtained using the T2*-weighted echo-planar sequence (TR=2000ms, TE=25ms, voxel size=3×3×3mm, flip angle=75 degrees, FoV=240mm, slice=40).

##### fMRI data preprocessing

The preprocessing pipeline consisted of three stages. The first stage utilized fMRIPrep version 20.2.1 (Esteban et al., 2019) for both structural and functional imaging preprocessing. For structural images, T1-weighted scans were corrected for intensity non-uniformity and skull-stripped, followed by tissue segmentation into gray matter, white matter, and cerebrospinal fluid. No cortical surface reconstruction was performed. The T1-weighted images were spatially normalized to the MNI152NLin2009cAsym template at 2 mm resolution using nonlinear registration. Functional preprocessing included the generation of a reference image, correction for head motion, and susceptibility distortion correction using a fieldmap-free approach based on nonlinear registration between the BOLD reference and the T1-weighted image. Functional images were then coregistered to the corresponding T1-weighted images and normalized to the same MNI space. Several confounding time series were estimated from the preprocessed BOLD data, including motion parameters, framewise displacement, DVARS, and physiological noise components derived from white matter and cerebrospinal fluid masks using component-based noise correction. Finally, spatial smoothing was applied using a Gaussian kernel with a 6 mm full-width-at-half-maximum (FWHM).

In the second stage, we fitted a general linear model (GLM) to each participant’s voxelwise time series. Nuisance regressors included an intercept, a linear trend, and 24 motion-related regressors comprising the six rigid-body realignment parameters, their squared terms, their first temporal differences, and the squared first temporal differences. In addition, spike regressors were identified directly from the BOLD time series using the find_spikes function in Nltools, with both the global-signal spike cutoff and frame-difference spike cutoff set to three standard deviations. These spike time points were modeled as nuisance regressors rather than removed or censored from the time series. No CSF, white-matter, CompCor, or whole-brain global signal regressor was included in the GLM. Framewise displacement (FD) and DVARS were estimated by fMRIPrep for quality-control purposes but were not used as threshold-based criteria for participant exclusion, volume censoring, or spike definition. No fMRI volumes were deleted or censored, and no additional temporal filtering (high-pass, low-pass, or band-pass filtering) was applied to the residual time series. The resulting residual time series were used in subsequent analyses. For each video, the analyzed time series included all TRs from the first TR corresponding to video onset through video offset. No temporal shift was applied to account for the hemodynamic lag, and no additional TRs were added or removed at either the beginning or end of the video to account for video transitions (L. J. Chang et al., 2021; P.-H. A. Chen et al., 2020). The third stage was to divide the whole brain into 100 non-overlapping regions of interest (ROIs) based on meta-analytic functional coactivation data from the Neurosynth database (de la Vega et al., 2016). Subsequently, the residual time series data in all voxels within each ROI were averaged, generating the neural dynamics for 100 ROIs.

#### B.3. Intersubject representational similarity analysis

We utilized ISRSA to test whether intersubject variation in national identity ideology was reflected in intersubject variation in neural dynamics during viewing of the pro-China video (Figure 1). For the national identity ideology, we calculated intersubject similarity matrices using item-level response patterns for Greater Chinese ideology and Taiwanese refinement ideology separately. Similarity between each pair of participants was computed using Pearson correlation across all items within each subscale, and these similarities were converted into similarity matrices. This operationalization indexed similarity in the configuration of item responses rather than proximity in participants’ mean level of endorsement. It was used to test whether participants with more similar patterns of responses across the Greater Chinese ideology items also exhibited more similar neural dynamics. Alternative analyses using absolute distance between mean scores and Euclidean distance between standardized item responses are reported in the Supplementary Materials. For neural dynamics, we computed intersubject similarity matrices separately for each of the 100 ROIs by computing Pearson correlations between each pair of participants’ ROI time series during the pro-China video. For each brain region, we then quantified the correspondence between the item-response profile similarity matrix and the neural similarity matrix using Spearman rank correlation. Statistical significance was evaluated using Mantel permutation tests (Mantel, 1967; Nummenmaa et al., 2012) with 10,000 permutations. For each permutation, participant labels of the neural similarity matrix were randomly permuted by jointly applying the same permutation to its rows and columns, thereby preserving the symmetry and dependency structure of the matrix while disrupting its correspondence with the ideology-similarity matrix. The Spearman correlation between the permuted neural similarity matrix and the fixed ideology-similarity matrix was then recomputed to generate the permutation null distribution. Because our directional hypothesis predicted that greater similarity in Greater Chinese ideology would be associated with greater neural similarity, statistical significance was assessed using the upper tail of the permutation distribution, corresponding to the alternative hypothesis that the ISRSA correlation was greater than zero (P.-H. A. Chen et al., 2020; Hsiao et al., 2024; Parkinson et al., 2018). Resulting p-values were corrected for multiple comparisons across ROIs using false discovery rate control, Benjamini–Hochberg, *q* < 0.05 (Benjamini & Hochberg, 1995), using the fdr function from Nltools (L. Chang et al., 2018).

**Figure 1.**
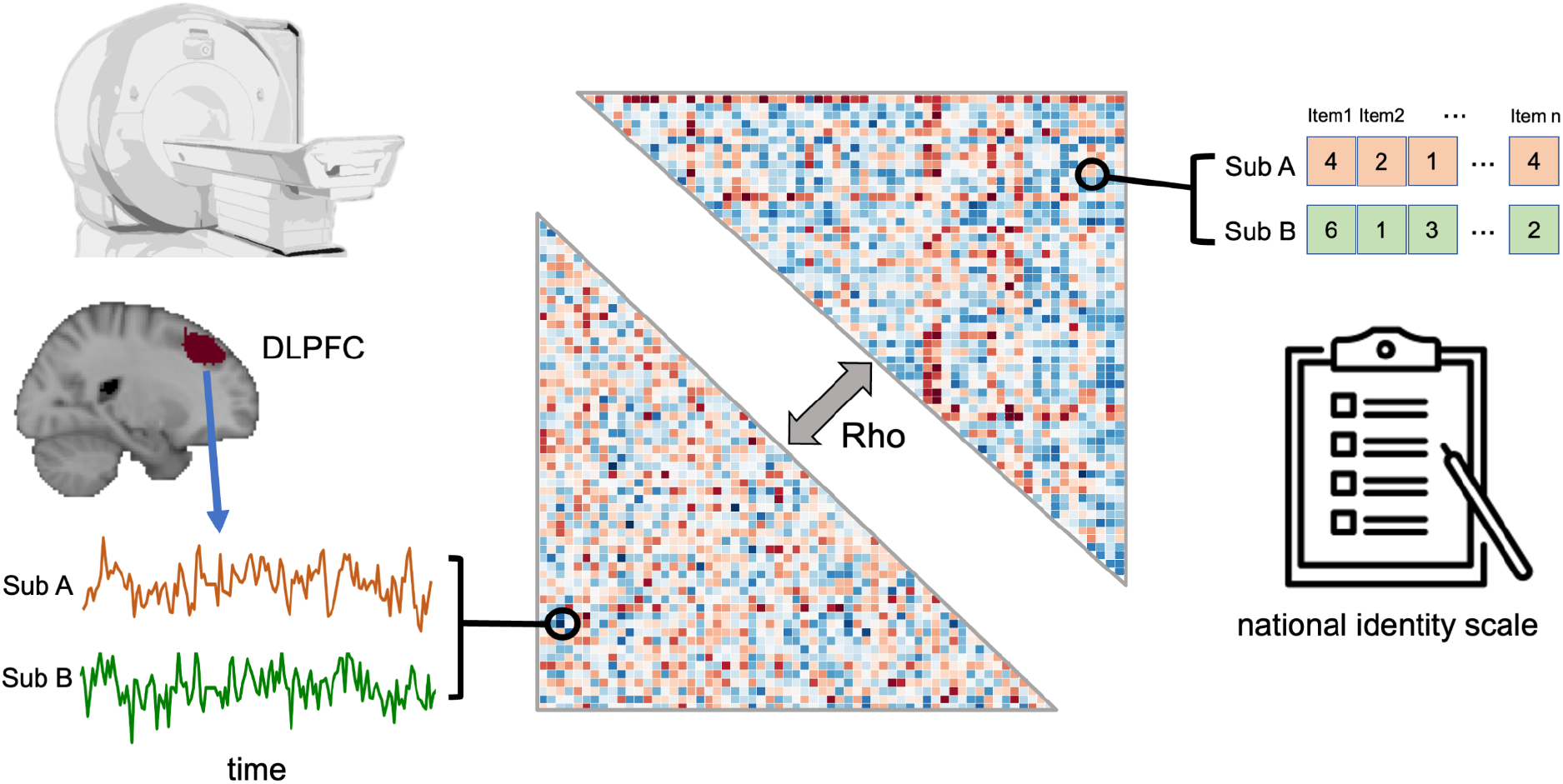
Intersubject representational similarity analysis overview. We computed intersubject similarity matrices from item-level responses for Greater Chinese ideology and Taiwanese refinement ideology using Pearson correlations between participants. We computed corresponding neural similarity matrices by correlating each pair of participants’ ROI time series for each of the 100 ROIs. For each ROI, we evaluated the correspondence between item-response profile similarity and neural dynamic similarity using Spearman rank correlation. Statistical significance was evaluated using Mantel permutation tests with 10,000 iterations, and results were corrected for multiple comparisons across ROIs using false discovery rate control at q less than 0.05.

To compare the associations across ideology dimensions, we repeated the ISRSA procedure for Taiwanese refinement ideology and directly compared the ISRSA coefficients for Greater Chinese ideology and Taiwanese refinement ideology within each ROI during the pro-China video. Statistical significance of the difference between the two dependent coefficients was evaluated using 10,000 participant-label permutations, with the same permutation applied to both ideology similarity matrices on each iteration to preserve their dependency. Permutation p values were calculated from the observed difference in Spearman correlations and corrected across the 100 ROIs using false discovery rate (FDR) correction. To compare the association across stimuli, we repeated the ISRSA procedure using neural dynamics during the 91-s nature control video and the additional 271-s non-political narrative control video, Farewell. We then directly compared the ISRSA coefficients between the pro-China video and each control video within each ROI. For each comparison, the pro-China and control-video time series were concatenated within each participant, and each participant’s concatenated time series was independently circularly shifted by a randomly selected temporal offset. The shifted time series were then divided back into the original pro-China and control-video segments, preserving the duration of each video. Neural intersubject similarity matrices and ISRSA coefficients were recomputed for both videos, and the difference between coefficients (Δ*ρ* = *ρ*_pro-China - *ρ*_control) was calculated. This procedure was repeated 10,000 times to generate a null distribution of Δ*ρ* while preserving the temporal autocorrelation of the time series. One-tailed permutation p values tested whether the ISRSA association was stronger during the pro-China video than during the corresponding control video, followed by FDR correction across the 100 ROIs.

#### B.4. Conducting the within-vs-between-group similarity analysis

To further characterize the ISRSA effects descriptively, we followed prior work using a within-vs-between-group similarity analysis (Broom et al., 2022; Leong et al., 2020; Zebarjadi et al., 2026). Participants were divided into relatively higher-scoring and relatively lower-scoring groups according to whether their Greater Chinese ideology scores were above or below the sample median. In Study 2 (N = 60), Greater Chinese ideology scores ranged from 1.18 to 4.82 (*M* = 2.72, *SD* = 0.83, median = 2.73), with 30 participants below and 30 above the median. The relatively lower-scoring group had a mean score of 2.05 (*SD* = 0.44, range = 1.18–2.64), whereas the relatively higher-scoring group had a mean score of 3.39 (*SD* = 0.56, range = 2.82–4.82). These labels therefore refer to participants’ relative position within the present sample rather than high or low absolute endorsement, and the score distribution is shown in Figure S1.

For each ROI and each participant, we computed within-group similarity as the average Fisher z-transformed correlation between that participant’s ROI time series and the time series of all other participants within the same group (Figure 2A). We computed between-group similarity in the same way using participants from the opposite group (Figure 2A). We then compared within group and between-group similarity using paired sample t tests (Figure 2B). Within each ideology group, statistical significance was evaluated using a paired permutation test with 10,000 iterations. For each permutation, the within-group and between-group similarity values were randomly exchanged for a subset of participants, thereby randomly reversing the sign of their within-minus-between difference scores. This procedure generated a null distribution of paired-samples t statistics for evaluating the observed within-versus-between similarity effect. We applied false discovery rate correction across regions with q less than 0.05.

**Figure 2.**
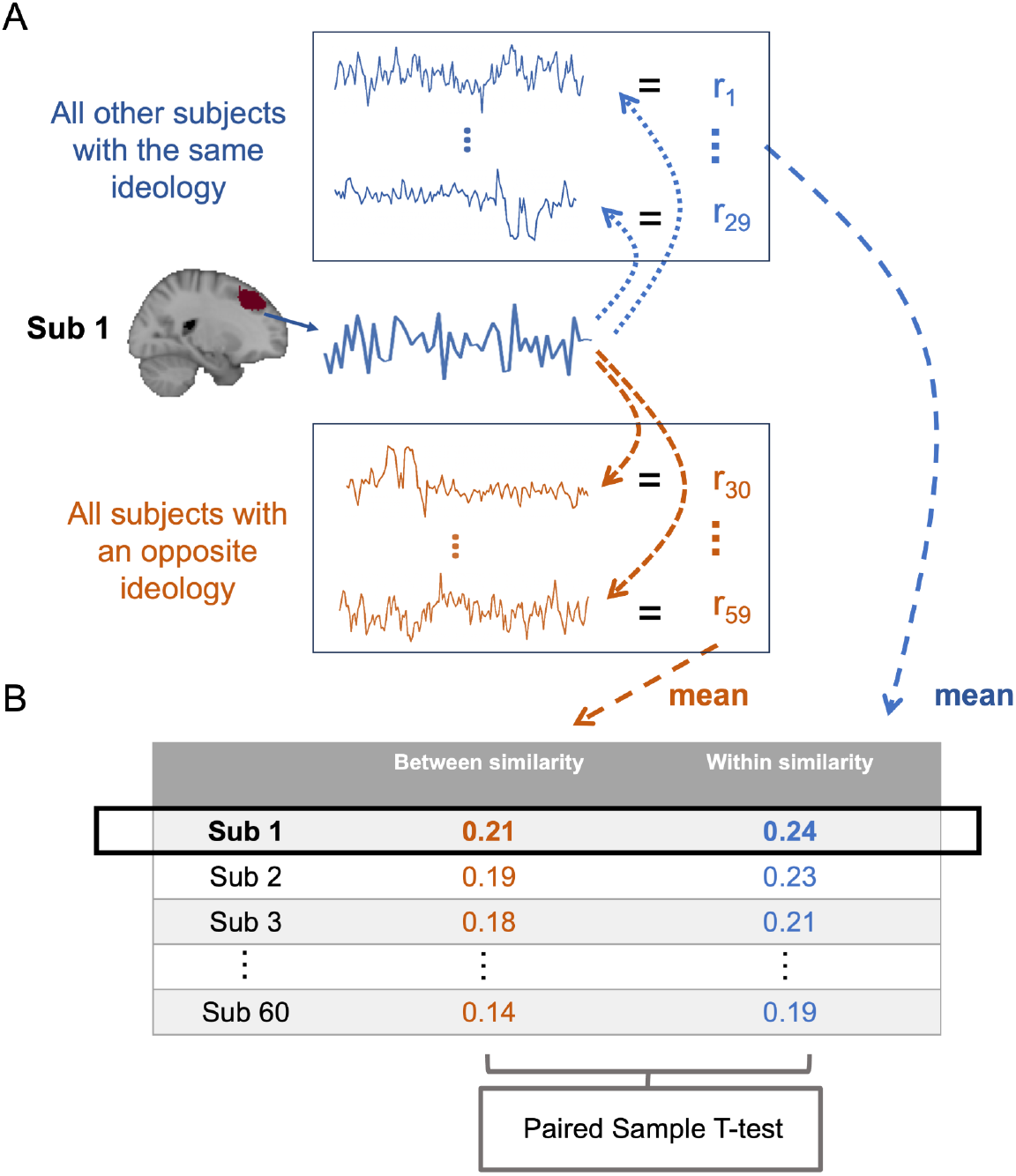
Within-vs-between-group neural similarity analysis. Participants were divided into relatively higher-scoring and relatively lower-scoring Greater Chinese ideology groups using a median split. For each participant and each ROI, we computed within-group similarity as the mean Fisher z-transformed Pearson correlation between that participant’s ROI time series and the time series of all other participants in the same group. We computed between-group similarity as the mean Pearson correlation between that participant’s time series and those of participants in the opposite group. Within group and between-group similarity were then compared using paired-sample t tests across participants for each ROI.

To directly test whether the within-versus-between similarity effect differed between the relatively lower- and relatively higher-scoring groups, we additionally compared the participant-level within-minus-between similarity scores between groups. Statistical significance was assessed using 10,000 permutations combining random reassignment of group labels with participant-level sign flipping, and one-tailed p-values were computed for the hypothesized direction of a larger effect in the relatively lower-scoring group. Resulting p-values were corrected across the 100 ROIs using FDR correction. To assess whether the findings depended on the median split, we repeated the analysis using alternative group definitions based on scores at least 0.25 SD and 0.5 SD below or above the sample mean, excluding participants whose scores fell between the corresponding thresholds.

## Results

### C. Study 1

We first tested whether individual differences in national identity ideology were associated with positive engagement with the pro-China video. In line with our hypothesis, Greater Chinese ideology was positively associated with positive engagement during the pro-China video, *r* = .47, 95% CI [.23, .66], *p* < .001 (Figure 3A). In contrast, Taiwanese refinement ideology was not significantly associated with positive engagement, *r* = −.18, 95% CI [−.43, .10], *p* = .20 (Figure 3B). To directly test whether engagement with the pro-China video was more strongly associated with Greater Chinese ideology than with Taiwanese refinement ideology, we compared the two dependent correlations using Williams’ test. The association was significantly stronger for Greater Chinese ideology, *t*(49) = 3.23, *p* = .002. Greater Chinese ideology was not significantly associated with positive engagement during the control video, *r* = .13, 95% CI [−.15, .39], *p* = .35. A direct comparison further showed that the association between Greater Chinese ideology and positive engagement was significantly stronger for the pro-China video than for the control video, Williams *t*(49) = 2.10, *p* = .041. Together, these direct comparisons showed that the association with positive engagement was stronger for Greater Chinese ideology than for Taiwanese refinement ideology and stronger for the pro-China video than for the control video.

**Figure 3.**
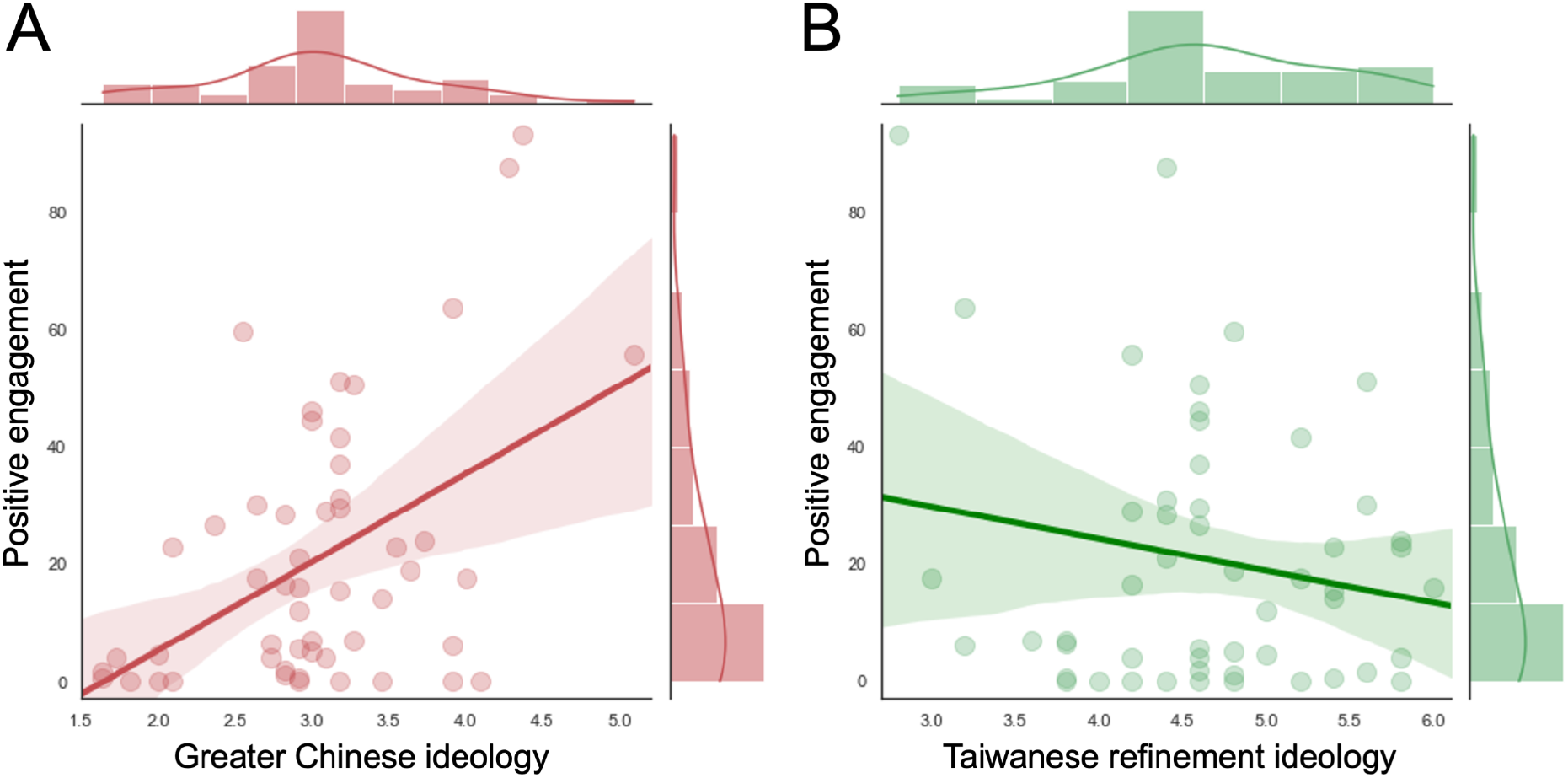
The association between national identity and positive engagement during viewing of the pro-China video. (A) Greater Chinese ideology was positively associated with positive engagement. (B) By contrast, the association between Taiwanese refinement ideology and positive engagement with the pro-China video was not significant.

### D. Study 2

Building directly on the behavioral findings from Study 1, Study 2 recruited an independent group of participants and examined whether individual differences in national identity ideology were reflected in neural dynamics during naturalistic viewing of the pro-China video. Participants viewed the same videos during fMRI and subsequently completed the same national identity ideology scale and comparable post-viewing ratings, although the emotion-rating items sightly differed between studies. Replicating our behavioral findings from Study 1, Greater Chinese ideology was associated with higher positive engagement of the pro-China video, *r* = 0.48, 95% CI [0.26, 0.65], *p* < .001, whereas Taiwanese refinement ideology showed no such association, *r* = −0.10, 95% CI [−0.34, 0.16], *p* = .465, supporting the reliability of the ideology–engagement association in an independent sample. Liking and resonance were again strongly correlated in Study 2, *r* = .85, 95% CI [.76, .91], *p* < .001, with a standardized two-item coefficient alpha of .92. Greater Chinese ideology was not associated with mean framewise displacement during the pro-China video, *r* = .013, *p* = .923, indicating that ideological differences were not systematically related to head motion.

#### D.1. ISRSA revealed significant associations between Greater Chinese ideology and neural dynamics in the frontoparietal and default mode networks

Using the item-response profile similarity matrix, we tested whether participants with more similar Greater Chinese ideology item-response profiles also showed more similar neural dynamics during viewing of the pro-China video. Our results showed that Greater Chinese ideology item-response profile similarity was associated with neural similarity within the frontoparietal network, including the dorsal anterior cingulate cortex (dACC), *r* = 0.13, *p* = 0.003; dorsolateral prefrontal cortex (DLPFC), *r* = 0.11, *p* = 0.002; pre-supplementary motor area (pre-SMA), *r* = 0.10, *p* = 0.004; and bilateral insula, *r* = 0.12, *p* < 0.001 (Figure 4A). We also found similar associations in regions within the default mode network, including the posterior cingulate cortex (PCC), *r* = 0.10, *p* = 0.006, and precuneus, *r* = 0.10, *p* = 0.003 (Figure 4A). The p-values reported here are uncorrected permutation p-values; all reported ROIs survived false discovery rate correction across the 100 ROIs at *q* < .05. Complete results for all 100 ROIs, including ROI labels, MNI centroid coordinates, effect sizes, uncorrected p-values, and FDR-adjusted q-values, are provided in Table S3. By contrast, no significant association between Taiwanese refinement ideology item-response profile similarity and neural similarity survived FDR correction across the 100 ROIs (Figure 4B). A direct permutation comparison further showed that the ISRSA association was significantly stronger for Greater Chinese ideology than for Taiwanese refinement ideology in the pre-supplementary motor area (Δ*ρ* = .156, *p* = .0027) and bilateral anterior insula (Δ*ρ* = .163, *p* = .0019), with both differences surviving FDR correction across the 100 ROIs (Figure S2). Lastly, we evaluated stimulus specificity by repeating the ISRSA for neural dynamics during the nature video. For the nature video, item-response profile similarity for neither Greater Chinese ideology nor Taiwanese refinement ideology was significantly associated with neural similarity after FDR correction.

**Figure 4.**
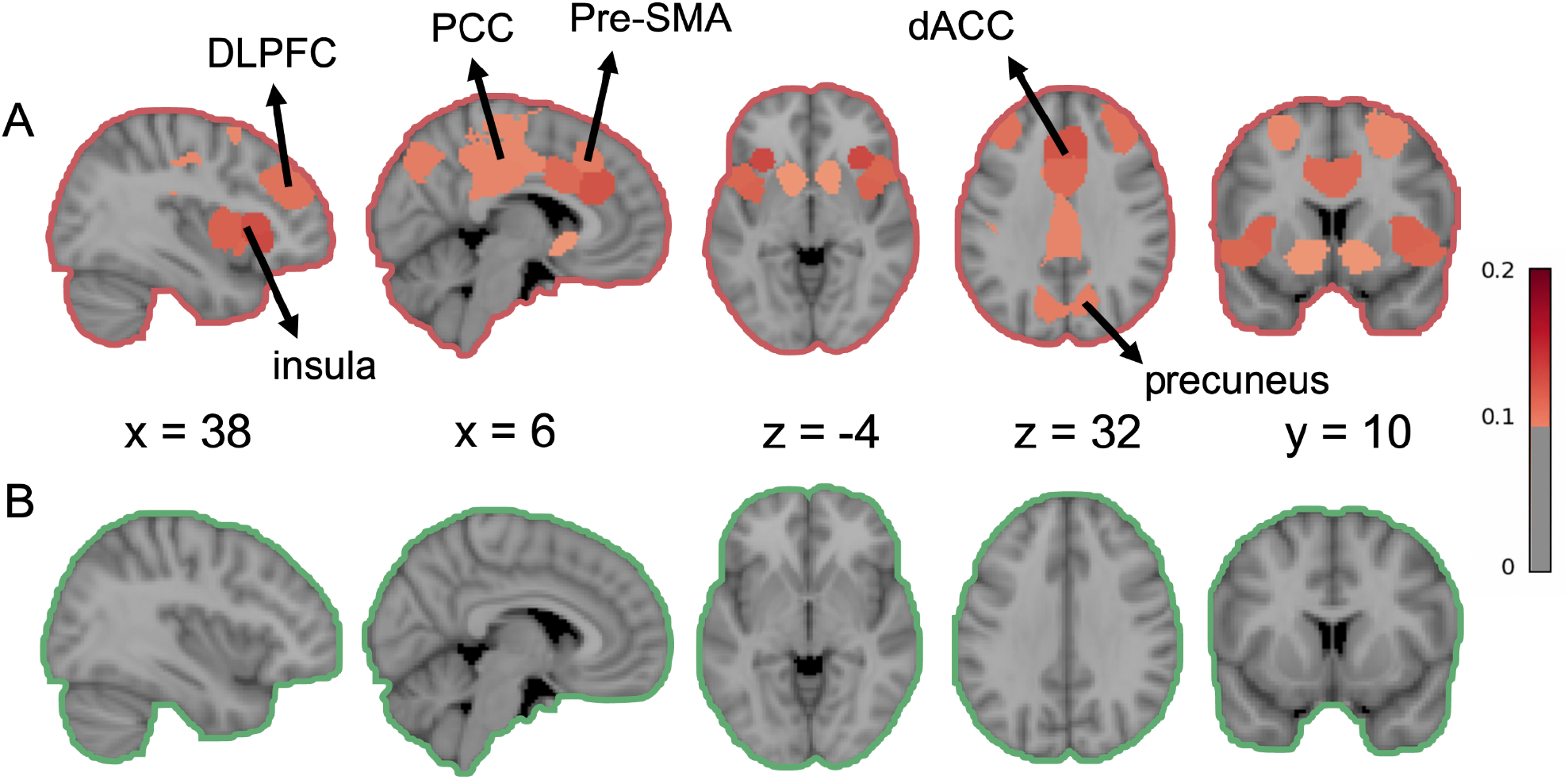
Intersubject representational similarity analysis (ISRSA) results. (A) Greater Chinese ideology item-response profile similarity was positively associated with neural similarity in the dorsal anterior cingulate cortex (dACC), dorsolateral prefrontal cortex (DLPFC), bilateral anterior insula, posterior cingulate cortex (PCC), precuneus, and pre-supplementary motor area (pre-SMA). (B) No significant associations were observed for Taiwanese refinement ideology. The color scale represents Spearman’ s *ρ*, with warmer colors indicating stronger positive associations. Only ROIs surviving false discovery rate (FDR) correction across the 100 ROIs (*q* < .05) are displayed in color, whereas nonsignificant ROIs are shown in gray.

Additional analyses were conducted to examine whether the association between Greater Chinese ideology item-response profile similarity and neural similarity differed between the pro-China video and two control videos and remained after accounting for alternative sources of inter-individual similarity. A direct permutation comparison showed that this association was significantly stronger during the pro-China video than during the nature control video in the bilateral precuneus (Δ*ρ* = .112, *p* = .0007), pre-supplementary motor area (Δ*ρ* = .100, *p* = .0019), bilateral anterior insula (Δ*ρ* = .094, *p* = .0025), bilateral dorsolateral prefrontal cortex (Δ*ρ* = .086, *p* = .0046), and dorsal anterior cingulate cortex (Δ*ρ* = .098, *p* = .0020). All five differences survived FDR correction across the 100 ROIs (Figure S3). To provide an additional comparison using a longer, non-political narrative control, we repeated the analysis using the 271-s Farewell video. The association was significantly stronger during the pro-China video than during the Farewell control video in the bilateral anterior insula (Δ*ρ* = .128, *p* = .0003), bilateral dorsolateral prefrontal cortex (Δ*ρ* = .083, *p* = .0069), and dorsal anterior cingulate cortex (Δ*ρ* = .088, *p* = .0041). All three differences survived FDR correction across the 100 ROIs (Figure S4).

We further examined whether the association could be accounted for by similarities in participants’ comprehension of the video or head motion. After controlling for pairwise similarity in comprehension and in the six rigid-body head-motion parameters, the association between Greater Chinese ideology item-response profile similarity and neural similarity remained significant in the bilateral precuneus (*β* = .066, *p* = .0036), bilateral posterior cingulate cortex (*β* = .057, *p* = .0055), pre-supplementary motor area (*β* = .064, *p* = .0029), bilateral dorsolateral prefrontal cortex (*β* = .062, *p* = .0042), dorsal anterior cingulate cortex (*β* = .079, *p* = .0052), and bilateral anterior insula (*β* = .068, *p* = .0008; Figure S5). Together, these additional analyses showed that the association was stronger during the pro-China video than during the two control videos in several regions and remained significant after accounting for similarities in comprehension and head motion.

#### D.2. Individuals with relatively lower Greater Chinese ideology showed greater within-vs-between-group similarity in the frontoparietal and default mode networks

The ISRSA results demonstrated that participants with more similar Greater Chinese ideology item-response profiles exhibited similar brain response patterns during the pro-China video. This profile-based association did not establish whether the pattern was expressed similarly across relatively lower-scoring and relatively higher-scoring portions of the sample. To further interpret the ISRSA effects, we conducted a within-vs-between-group similarity analysis by dividing participants into relatively higher-scoring and relatively lower-scoring Greater Chinese ideology groups using a median split. Across 100 ROIs, within-group similarity exceeded between-group similarity primarily for participants from the relatively lower-scoring group. This pattern was observed in the DLPFC, *t* = 6.17, *p* = .002; pre-SMA, *t* = 7.16, *p* < .001; PCC, *t* = 5.76, *p* = .001; and precuneus, *t* = 5.95, *p* = .001 (Figure 5A). By contrast, the relatively higher-scoring group showed no such differences in the above ROIs (Figure 5B). These findings suggest that the profile-related neural similarity was observed primarily among participants with relatively lower Greater Chinese ideology scores. A direct comparison further showed that the within-versus-between similarity effect was significantly greater in the relatively lower-scoring than in the relatively higher-scoring group in the pre-SMA, *t* = 4.968, *p* = .0024; bilateral DLPFC, *t* = 5.908, *p* = .0003; and bilateral anterior insula, *t* = 5.067, *p* = .0017. All three effects survived FDR correction across the 100 ROIs (Figure S6). To assess whether this pattern depended on the median split, we repeated the analysis using more stringent group definitions. Using thresholds of ±0.25 SD from the sample mean, significant group differences were observed in the bilateral precuneus, PCC, pre-SMA, DLPFC, dACC, and bilateral anterior insula, with all effects surviving FDR correction across the 100 ROIs (Figure S7). Using thresholds of ±0.5 SD from the sample mean, significant group differences remained in the bilateral precuneus, PCC, pre-SMA, and DLPFC, again surviving FDR correction across the 100 ROIs (Figure S8). These sensitivity analyses indicate that the overall pattern was not specific to the median-split threshold.

**Figure 5.**
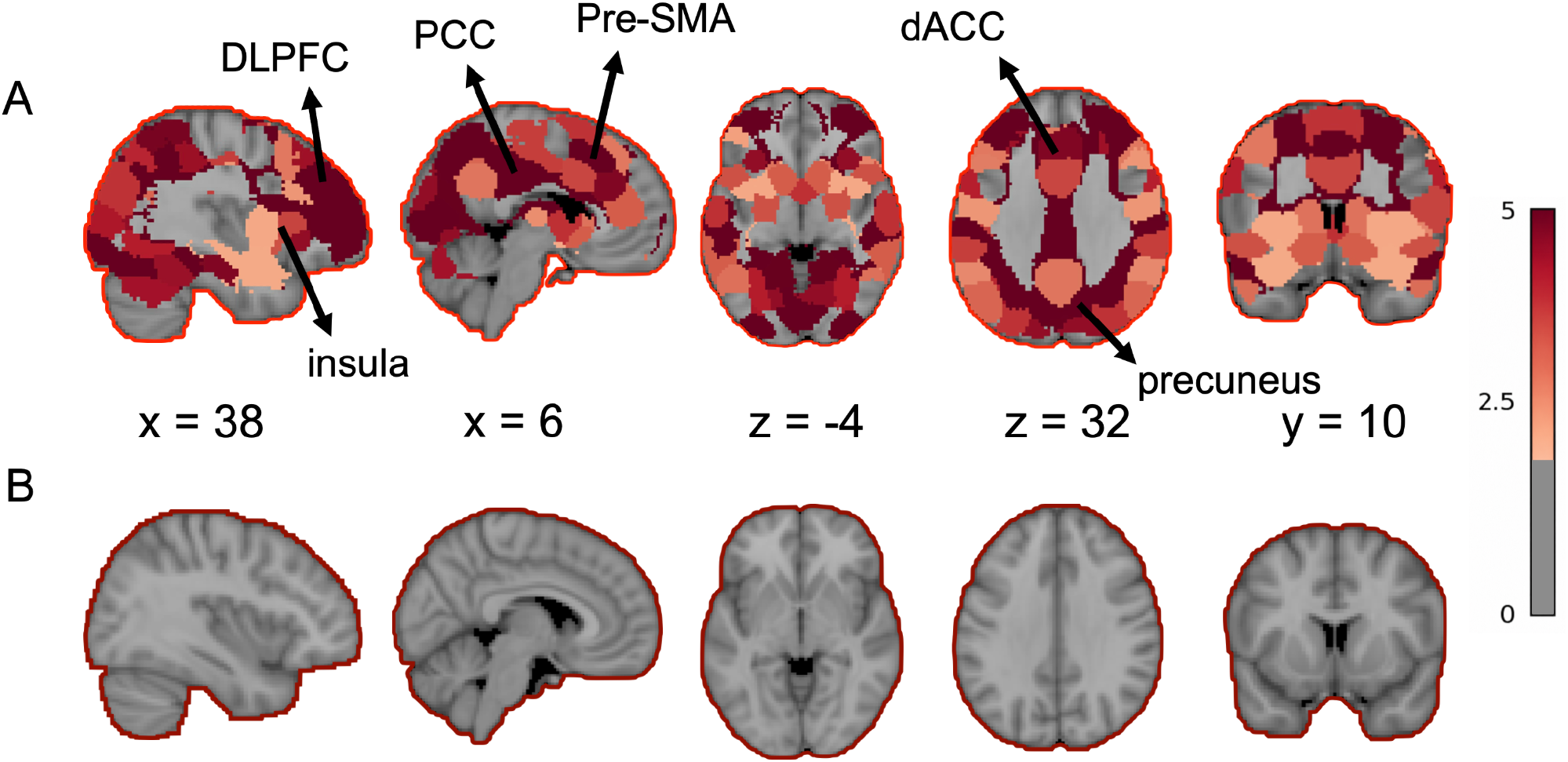
Within-group versus between-group neural similarity analysis. (A) Participants with relatively lower Greater Chinese ideology showed significantly greater within-group than between-group neural similarity in the dorsolateral prefrontal cortex (DLPFC), posterior cingulate cortex (PCC), precuneus, and pre-supplementary motor area (pre-SMA). (B) No significant within-group versus between-group differences were observed among participants with relatively higher Greater Chinese ideology. The color scale represents *t* values for the within-group versus between-group neural similarity comparison, with warmer colors indicating stronger within-group relative to between-group similarity. Only ROIs surviving false discovery rate (FDR) correction across the 100 ROIs (*q* < .05) are displayed in color, whereas ROIs not surviving this threshold are shown in gray.

We also conducted the same within-versus-betweengroup analysis using Taiwanese refinement ideology to examine whether this alternative dimension of national identity ideology showed a similar subgroup pattern. Using a strict median split (median = 4.60), 26 participants were relatively higher-scoring, 29 were relatively lower-scoring, and five participants who scored exactly at the median were excluded. No ROI showed a significant within-group versus between-group difference after FDR correction across the 100 ROIs (minimum *q* values = .494, .496, and .518 for the relatively lower-scoring, relatively higher-scoring, and pooled analyses, respectively). Thus, the asymmetric within-group similarity pattern was not observed for Taiwanese refinement ideology.

## Discussion

This study examined how national identity ideology relates to subjective experience and brain dynamics during naturalistic viewing of a pro-China video. Across two independent studies, we found that individuals with higher Greater Chinese ideology reported having greater positive engagement with the pro-China video, indicating that ideological alignment was expressed as stronger positive engagement with ideologically congruent content (Katabi et al., 2023; Mattingly & Yao, 2022; You, 2020). In Study 2, participants with more similar Greater Chinese ideology item-response profiles also showed more similar neural dynamics during viewing, as quantified by intersubject representational similarity analysis. This finding was selective to Greater Chinese ideology and the pro-China narrative, and was not observed for Taiwanese refinement ideology or during viewing of the control video. Together, the findings indicate that Greater Chinese ideology is associated with subjective engagement and neural similarity during a pro-China narrative, while the psychological processes underlying these associations remain uncertain.

Similarity in neural dynamics associated with Greater Chinese ideology was primarily observed in brain systems involved in cognitive control and social information processing. These included regions associated with the frontoparietal control and salience networks, including the DLPFC, pre-SMA, dACC, and anterior insula, as well as regions within the default mode network, including the PCC and precuneus. Prior work has shown that the frontoparietal control network supports evaluation, conflict monitoring, and regulation when individuals process information that is in line with their internal goals or beliefs (Cole et al., 2013; Harding et al., 2015; Marek & Dosenbach, 2018). In the present study, similarity in neural dynamics within these regions may reflect comparable evaluative, conflict-monitoring, or regulatory processes among participants with more similar Greater Chinese ideology item-response profiles as they watched the narrative, particularly when assessing its relevance to their own beliefs. However, similar neural dynamics do not necessarily indicate that participants engaged in the same psychological processes. This interpretation remains speculative, as these processes were not directly measured or experimentally tested in the present study. At the same time, similarity in neural dynamics within the PCC and precuneus is consistent with their well-established role in constructing higher-level meaning, interpreting social information, and representing group identity (Menon, 2023; Song et al., 2021; Vollberg & Cikara, 2018; E. Yang et al., 2023). Although we did not directly measure participants’ interpretations of the narrative, the similarity in PCC and precuneus dynamics among participants with more similar ideologies may be consistent with similar higher-level processing of social and identity-related information. However, this possibility remains tentative and should be tested using direct measures of narrative interpretation in future studies. Taken together, these results indicate that ideological similarity was associated with similarity in neural dynamics within regions previously implicated in evaluation, cognitive control, and social information processing. The specific psychological processes underlying this neural similarity, including whether it reflects similar interpretations of the narrative, remain to be directly examined in future studies.

Expanding on the above result, our study further found that intersubject variations in national identity ideology were associated with differences in neural similarity across relatively lower- and higher-scoring participants while viewing an identity-related video in Taiwan, where national identity is actively contested and frequently invoked in public discourse (Chu, 2004; L.-L. Huang, 2007; Wu, 2005; Zhong, 2016). In such settings, narratives about national belonging are likely to prompt active interpretation and evaluation rather than passive viewing. Within this context, similarity in neural dynamics associated with Greater Chinese ideology was not evenly distributed across the relatively lower- and higher-scoring portions of the present sample. Instead, direct group comparisons showed that the within-versus-between neural similarity effect was stronger among participants with relatively lower Greater Chinese ideology scores. This pattern differed from some previous findings related to political partisanship, where neural similarity has been observed at both ideological extremes (de Bruin et al., 2023; Leong et al., 2020; van Baar & FeldmanHall, 2022; Zmigrod & Tsakiris, 2021). However, recent work has also reported asymmetry. Zebarjadi et al. (2026) found a larger difference between within- and between-group neural similarity in the DLPFC among less immigration-supportive participants during anti-immigration narratives. Their findings further suggest that differences in neural responses to political narratives can emerge even among participants with relatively small differences in self-reported political perspectives. Together, these findings suggest that the relationship between ideology and neural similarity may vary across individuals and narrative contexts, rather than following the same pattern across the ideological continuum. These results suggest that sharing similar ideological views does not always lead to similar patterns of neural processing. Rather, the association between national identity ideology and neural dynamics may depend on how individuals engage with a given narrative, raising the possibility that the association between ideology and neural dynamics differs across relatively lower- and higher-scoring portions of a sample (Castelli & Carraro, 2011; Jost, 2017; Jost et al., 2022; Jost & Amodio, 2012; Kahan, 2013; Katabi et al., 2023).

A possible interpretation of this difference between relatively lower- and higher-scoring participants is that individuals with relatively lower Greater Chinese ideology scores may have engaged with the pro-China narrative differently than those with relatively higher scores. Participants with relatively lower scores may have perceived the narrative as less aligned with their existing views and engaged in greater evaluation of its content (Kunda, 1990; Lord et al., 1979). However, lower endorsement of Greater Chinese ideology does not necessarily indicate that participants experienced the narrative as inconsistent with their beliefs or perceived it as threatening. Previous studies have shown that processing belief-incongruent information increases cognitive effort and recruits the frontoparietal control network (Dixon et al., 2018; Izuma et al., 2010; van Veen et al., 2009). The involvement of regions associated with the frontoparietal control and salience networks is compatible with the possibility that participants evaluated the narrative in relation to their existing beliefs (Cole et al., 2013; Dosenbach et al., 2008). Similarity in neural dynamics within these regions may reflect comparable evaluative demands, but it does not establish that participants experienced greater conflict, cognitive effort, or active regulation (Braun et al., 2025; Németh et al., 2024). Additional analyses of the available behavioral ratings provide further context for this interpretation. Across both studies, Greater Chinese ideology was positively associated with resonance, while willingness to share also showed a positive association that remained significant after correction for multiple comparisons in Study 2. In contrast, Greater Chinese ideology was not significantly associated with negative emotion in Study 1 or comprehension in either study. These findings provide additional evidence that Greater Chinese ideology was related to participants’ subjective engagement with the narrative. However, these ratings do not directly establish whether participants experienced belief incongruence, ideological threat, or greater regulatory demands. Furthermore, Taiwanese refinement ideology did not show comparable associations with either positive engagement or neural dynamics, and the same analyses applied to the nature video also revealed no ideology-related neural similarity. These findings provide some evidence that the observed effects were specific to Greater Chinese ideology during the pro-China narrative, although they do not by themselves identify the psychological process underlying the neural similarity.

Recent studies have examined how the brain responds to challenges to existing beliefs and how neural responses relate to changes in narrative interpretation. Kossowska et al. (2026) investigated neural responses to counterarguments challenging political and non-political beliefs, revealing distinct patterns of activity in the DMN and cognitive control regions, particularly the DLPFC and pre-SMA. Importantly, their study directly presented belief-challenging information and assessed subsequent belief change, whereas the present study did not manipulate narrative–belief congruence or directly measure perceived conflict. Boiman et al. (2026) further found that longitudinal changes in neural responses to political videos were associated with changes in narrative interpretation and political in-group affiliation, whereas associations with changes in ideological positions were not statistically significant. These findings highlight the importance of distinguishing ideological beliefs from group identification and narrative interpretation, which may contribute differently to neural responses during political communication. Together, these studies suggest that neural responses to political narratives may reflect several related psychological processes, while the present findings cannot determine which processes contributed to the observed ideology-related neural similarity.

Although the present findings provide evidence for an association between Greater Chinese ideology and neural similarity, several limitations should be considered when interpreting these results. First, participants in the present study were primarily from younger generations, but previous studies have shown that patterns of national identity ideology in Taiwan differ markedly across generations (G. A. Chang & Wang, 2005; C. Huang, 2019). Future studies should include participants from different generations and examine whether our current findings also extend to generations with different historical experiences in Taiwan. Second, the present study examined neural responses to a single political narrative and focused on one specific dimension of national identity ideology. While this design allowed researchers to isolate a theoretically meaningful ideological construct, it does not address whether the observed patterns generalize across different types of political narratives or other dimensions of national identity ideology. Future studies should vary narrative content and include additional dimensions of national identity ideology and examine whether ideological differences could reliably give rise to shared neural dynamics. Third, we did not directly measure how participants interpreted the narrative during viewing, whether they perceived it as inconsistent with their beliefs or threatening, or whether viewing elicited conflict, cognitive effort, or active regulation. We also did not independently assess political in-group affiliation. Although the available post-viewing ratings provided additional information about participants’ subjective engagement, they could not directly establish these psychological processes. The observed neural similarity therefore cannot distinguish shared narrative interpretation from other processes that may contribute to similar neural dynamics. Future studies could directly manipulate narrative–belief congruence and incorporate time-resolved measures of narrative interpretation, perceived threat, cognitive conflict, and group identification to better understand the psychological processes underlying ideology-related neural similarity. Fourth, the primary ISRSA quantified similarity in participants’ item-response profiles rather than proximity in their overall level of Greater Chinese ideology. The neural findings should therefore be interpreted as reflecting similarity in patterns of item responses rather than similarity in the overall strength of ideological endorsement. Fifth, no formal a priori sample-size calculation was conducted for either study. However, the sample sizes were comparable to those used in previous studies employing ISC and related intersubject neural similarity analyses (Baek et al., 2022; Nguyen et al., 2019; Parkinson et al., 2018; van Baar et al., 2021).

To conclude, this study shows that Greater Chinese ideology, one dimension of national identity ideology, is associated with how individuals experience and respond to the pro-China video examined here, at both the behavioral and neural levels. By focusing on Greater Chinese ideology in Taiwan, this work extends prior research beyond political partisan contexts and highlights national identity as a central ideological dimension. Across two independent studies, we found that higher Greater Chinese ideology scores were associated with greater positive engagement with this pro-China video. In the fMRI study, similarity in Greater Chinese ideology item-response profiles was associated with similarity in neural dynamics, particularly in brain networks involved in monitoring, evaluation, and social information processing. Together, these findings indicate that Greater Chinese ideology is associated with subjective engagement and neural similarity during pro-China narrative viewing, while the psychological processes underlying these associations remain to be tested directly.

## Notes

### Competing Interest Statement

The authors have declared no competing interest.

### Summary of Updates

The title, abstract, and multiple parts of the manuscript has been revised.

## References

Baek, E. C., Hyon, R., López, K., Finn, E. S., Porter, M. A., & Parkinson, C. (2022). In-degree centrality in a social network is linked to coordinated neural activity. Nature Communications, 13(1), 1118.

Benjamini, Y., & Hochberg, Y. (1995). Controlling the False Discovery Rate: A Practical and Powerful Approach to Multiple Testing. Journal of the Royal Statistical Society. Series B, Statistical Methodology, 57(1), 289–300.

Boiman, G., Ohad, T., Zvi, Y., Katabi, N., & Yeshurun, Y. (2026). Changes in political attitudes are associated with changes in neural responses to political content. Communications Psychology, 4(1), 29.

Braun, G., Yeshurun, Y., & Shetreet, E. (2025). Shared disbelief and shared belief: Belief and disbelief as drivers of interpersonal neural synchronization during narrative processing. Proceedings of the National Academy of Sciences of the United States of America, 122(23), e2422396122.

Broom, T. W., Stahl, J. L., Ping, E. E. C., & Wagner, D. D. (2022). They Saw a Debate: Political Polarization Is Associated with Greater Multivariate Neural Synchrony When Viewing the Opposing Candidate Speak. Journal of Cognitive Neuroscience, 35(1), 60–73.

Castelli, L., & Carraro, L. (2011). Ideology is related to basic cognitive processes involved in attitude formation. Journal of Experimental Social Psychology, 47(5), 1013–1016.

Chang, G. A., & Wang, Y. T. (2005). Taiwanese or Chinese? Independence or unification? An analysis of generational differences in Taiwan. Journal of Asian and African Studies. 10.1177/0021909605052938

Chang, L. J., Jolly, E., Cheong, J. H., Rapuano, K. M., Greenstein, N., Chen, P.-H. A., & Manning, J. R. (2021). Endogenous variation in ventromedial prefrontal cortex state dynamics during naturalistic viewing reflects affective experience. Science Advances, 7(17), DOI: 10.1126/sciadv.abf7129.

Chang, L., Jolly, E., Cheong, J. H., Burnashev, A., & Chen, A. (2018). cosanlab/nltools: 0.3.11. 10.5281/zenodo.2229813

Chen, P.-H. A., Jolly, E., Cheong, J. H., & Chang, L. J. (2020). Intersubject representational similarity analysis reveals individual variations in affective experience when watching erotic movies. NeuroImage, 216(116851), 116851.

Chen, P.-H. A., & Qu, Y. (2021). Taking a computational cultural neuroscience approach to study parent-child similarities in diverse cultural contexts. Frontiers in Human Neuroscience, 15, 703999.

Chen, W.-L., Lin, M.-J., & Yang, T.-T. (2023). Curriculum and national identity: Evidence from the 1997 curriculum reform in Taiwan. Journal of Development Economics, 163(103078), 103078.

Chou, F.-C. B., & Chen, P.-H. A. (2025). Beyond dyadic interaction and shared experience: Rethinking social connections. In Psychology of Learning and Motivation. Elsevier.

Chou, F.-C. B., Chen, Y.-C. C., Kuo, Y.-S. A., Hsiao, P.-Y. A., Lin, C. R., Huang, Y.-W. Y., Peng, W.-L., Pan, H.-J., & Chen, P.-H. A. (2025). Facial insights: An exploration of similarity in spontaneous facial behaviors during pandemic lockdown. Technology, Mind, and Behavior, 6(2), 212–224.

Chu, Y.-H. (2004). Taiwan’s national identity politics and the prospect of cross-strait relations. Asian Survey, 44(4), 484–512.

Cole, M. W., Reynolds, J. R., Power, J. D., Repovs, G., Anticevic, A., & Braver, T. S. (2013). Multi-task connectivity reveals flexible hubs for adaptive task control. Nature Neuroscience, 16(9), 1348–1355.

de Bruin, D., & FeldmanHall, O. (2025). Politically extreme individuals exhibit similar neural processing despite ideological differences. Journal of Personality and Social Psychology, 129(5), 816–833.

de Bruin, D., van Baar, J. M., Rodríguez, P. L., & FeldmanHall, O. (2023). Shared neural representations and temporal segmentation of political content predict ideological similarity. Science Advances, 9(5), eabq5920.

de la Vega, A., Chang, L. J., Banich, M. T., Wager, T. D., & Yarkoni, T. (2016). Large-Scale Meta-Analysis of Human Medial Frontal Cortex Reveals Tripartite Functional Organization. The Journal of Neuroscience: The Official Journal of the Society for Neuroscience, 36(24), 6553–6562.

Dixon, M. L., De La Vega, A., Mills, C., Andrews-Hanna, J., Spreng, R. N., Cole, M. W., & Christoff, K. (2018). Heterogeneity within the frontoparietal control network and its relationship to the default and dorsal attention networks. Proceedings of the National Academy of Sciences of the United States of America, 115(7), E1598–E1607.

Dosenbach, N. U. F., Fair, D. A., Cohen, A. L., Schlaggar, B. L., & Petersen, S. E. (2008). A dual-networks architecture of top-down control. Trends in Cognitive Sciences, 12(3), 99–105.

Esteban, O., Markiewicz, C. J., Blair, R. W., Moodie, C. A., Isik, A. I., Erramuzpe, A., Kent, J. D., Goncalves, M., DuPre, E., Snyder, M., Oya, H., Ghosh, S. S., Wright, J., Durnez, J., Poldrack, R. A., & Gorgolewski, K. J. (2019). fMRIPrep: a robust preprocessing pipeline for functional MRI. Nature Methods, 16(1), 111–116.

Finn, E. S., Glerean, E., Khojandi, A. Y., Nielson, D., Molfese, P. J., Handwerker, D. A., & Bandettini, P. A. (2020). Idiosynchrony: From shared responses to individual differences during naturalistic neuroimaging. NeuroImage, 215, 116828.

Fournier, P., Soroka, S., & Nir, L. (2020). Negativity biases and political ideology: A comparative test across 17 countries. The American Political Science Review, 114(3), 775–791.

Friedman, N. P., & Robbins, T. W. (2022). The role of prefrontal cortex in cognitive control and executive function. Neuropsychopharmacology: Official Publication of the American College of Neuropsychopharmacology, 47(1), 72–89.

Hakonen, M., Ikäheimonen, A., Hultèn, A., Kauttonen, J., Koskinen, M., Lin, F.-H., Lowe, A., Sams, M., & Jääskeläinen, I. P. (2022). Processing of an audiobook in the human brain is shaped by cultural family background. Brain Sciences, 12(5), 649.

Harding, I. H., Yücel, M., Harrison, B. J., Pantelis, C., & Breakspear, M. (2015). Effective connectivity within the frontoparietal control network differentiates cognitive control and working memory. NeuroImage, 106, 144–153.

Hsiao, P.-Y. A., Kim, M. J., Chou, F.-C. B., & Chen, P.-H. A. (2024). Intersubject representational similarity analysis uncovers the impact of state anxiety on brain activation patterns in the human extrastriate cortex. Brain Imaging and Behavior, 18(2), 1–9.

Huang, C. (2019). Generation effects? Evolution of independence–unification views in Taiwan, 1996–2016. Electoral Studies, 58, 103–112.

Huang, L.-L. (2007). M shape vs. Bell shape: The ideology of national identity and its psychological basis in Taiwan. Chinese Journal of Psychology, 49(4), 451–470. 10.6129/CJP.2007.4904.08

Huang, L.-L., Liu, J. H., & Chang, M. (2004). ‘The double identity’ of Taiwanese Chinese: A dilemma of politics and culture rooted in history. Asian Journal of Social Psychology, 7(2), 149–168.

Huber, R. E., Klucharev, V., & Rieskamp, J. (2015). Neural correlates of informational cascades: brain mechanisms of social influence on belief updating. Social Cognitive and Affective Neuroscience, 10(4), 589–597.

Izuma, K., Matsumoto, M., Murayama, K., Samejima, K., Sadato, N., & Matsumoto, K. (2010). Neural correlates of cognitive dissonance and choice-induced preference change. Proceedings of the National Academy of Sciences of the United States of America, 107(51), 22014–22019.

Jacobs, J. B. (2013). Whither taiwanization? The colonization, democratization and taiwanization of Taiwan. Japanese Journal of Political Science, 14(4), 567–586.

Jenke, L., & Huettel, S. A. (2016). Issues or identity? Cognitive foundations of voter choice. Trends in Cognitive Sciences, 20(11), 794–804.

Jost, J. T. (2017). Ideological asymmetries and the essence of political psychology: Presidential address. Political Psychology, 38(2), 167–208.

Jost, J. T., & Amodio, D. M. (2012). Political ideology as motivated social cognition: Behavioral and neuroscientific evidence. Motivation and Emotion, 36(1), 55–64.

Jost, J. T., Baldassarri, D. S., & Druckman, J. N. (2022). Cognitive–motivational mechanisms of political polarization in social-communicative contexts. Nature Reviews Psychology, 1(10), 560–576.

Jost, J. T., Nosek, B. A., & Gosling, S. D. (2008). Ideology: Its resurgence in social, personality, and political psychology. Perspectives on Psychological Science: A Journal of the Association for Psychological Science, 3(2), 126–136.

Kahan, D. M. (2013). Ideology, motivated reasoning, and cognitive reflection. Judgment and Decision Making, 8(4), 407–424.

Kaplan, J. T., Gimbel, S. I., & Harris, S. (2016). Neural correlates of maintaining one’s political beliefs in the face of counterevidence. Scientific Reports, 6(1). 10.1038/srep39589

Katabi, N., Simon, H., Yakim, S., Ravreby, I., Ohad, T., & Yeshurun, Y. (2023). Deeper Than You Think: Partisanship-Dependent Brain Responses in Early Sensory and Motor Brain Regions. The Journal of Neuroscience: The Official Journal of the Society for Neuroscience, 43(6), 1027–1037.

Knight, K. (2006). Transformations of the concept of ideology in the twentieth century. The American Political Science Review, 100(04), 619.

Kossowska, M., Szwed, P., Czarnek, G., Honcharowa, S., Domagalik, A., & Doradzińska, Ł. (2026). Neural correlates of belief change in political and non-political domains among left-wing individuals confronted with counterarguments. Scientific Reports, 16(1), 4895.

Kriegeskorte, N., & Douglas, P. K. (2018). Cognitive computational neuroscience. Nature Neuroscience, 21(9), 1148–1160.

Kriegeskorte, N., Mur, M., & Bandettini, P. (2008). Representational similarity analysis - connecting the branches of systems neuroscience. Frontiers in Systems Neuroscience, 2, 4.

Kunda, Z. (1990). The case for motivated reasoning. Psychological Bulletin, 108(3), 480–498.

Kunze, R. (2024). Homey foods: Domesticating memories of the martial-law era in Taiwan’s heritage tourism. Memory Studies, 17(1), 71–85.

Leong, Y. C., Chen, J., Willer, R., & Zaki, J. (2020). Conservative and liberal attitudes drive polarized neural responses to political content. Proceedings of the National Academy of Sciences of the United States of America, 117(44), 27731–27739.

Lord, C. G., Ross, L., & Lepper, M. R. (1979). Biased assimilation and attitude polarization: The effects of prior theories on subsequently considered evidence. Journal of Personality and Social Psychology, 37(11), 2098–2109.

Lyu, Z., & Zhou, H. (2023). Contesting master narratives: Renderings of national history by mainland China and Taiwan. The China Quarterly, 255, 768–784.

Mader, M., & Schoen, H. (2023). Stability of national-identity content: Level, predictors, and implications. Political Psychology, 44(4), 871–891.

Mantel, N. (1967). The detection of disease clustering and a generalized regression approach. Cancer Research, 27(2), 209–220.

Marek, S., & Dosenbach, N. U. F. (2018). The frontoparietal network: function, electrophysiology, and importance of individual precision mapping. Dialogues in Clinical Neuroscience, 20(2), 133–140.

Mattingly, D. C., & Yao, E. (2022). How soft propaganda persuades. Comparative Political Studies, 55(9), 1569–1594.

Meeus, J., Duriez, B., Vanbeselaere, N., & Boen, F. (2010). The role of national identity representation in the relation between in-group identification and out-group derogation: ethnic versus civic representation. The British Journal of Social Psychology, 49(Pt 2), 305–320.

Menon, V. (2023). 20 years of the default mode network: A review and synthesis. Neuron, 111(16), 2469–2487.

Mills, M., Smith, K. B., Hibbing, J. R., & Dodd, M. D. (2015). Obama cares about visuo-spatial attention: perception of political figures moves attention and determines gaze direction. Behavioural Brain Research, 278, 221–225.

Németh, D., Vékony, T., Orosz, G., Sarnyai, Z., & Zmigrod, L. (2024). The interplay between subcortical and prefrontal brain structures in shaping ideological belief formation and updating. Current Opinion in Behavioral Sciences, 57(101385), 101385.

Nguyen, M., Vanderwal, T., & Hasson, U. (2019). Shared understanding of narratives is correlated with shared neural responses. NeuroImage, 184, 161–170.

Nummenmaa, L., Glerean, E., Viinikainen, M., Jääskeläinen, I. P., Hari, R., & Sams, M. (2012). Emotions promote social interaction by synchronizing brain activity across individuals. Proceedings of the National Academy of Sciences of the United States of America, 109(24), 9599–9604.

Panish, A., & Nam, H. H. (2023). The neurobiology of political ideology: Theories, findings, and future directions. Social and Personality Psychology Compass. 10.1111/spc3.12916

Parkinson, C., Kleinbaum, A. M., & Wheatley, T. (2018). Similar neural responses predict friendship. Nature Communications, 9(1), 332.

Schreiber, D., Fonzo, G., Simmons, A. N., Dawes, C. T., Flagan, T., Fowler, J. H., & Paulus, M. P. (2013). Red brain, blue brain: evaluative processes differ in Democrats and Republicans. PloS One, 8(2), e52970.

Schulreich, S., & Schwabe, L. (2021). Causal role of the dorsolateral prefrontal cortex in belief updating under uncertainty. Cerebral Cortex (New York, N.Y.: 1991), 31(1), 184–200.

Shteynberg, G., Bramlett, J. M., Fles, E. H., & Cameron, J. (2016). The broadcast of shared attention and its impact on political persuasion. Journal of Personality and Social Psychology, 111(5), 665–673.

Sievers, B., Welker, C., Hasson, U., Kleinbaum, A. M., & Wheatley, T. (2024). Consensus-building conversation leads to neural alignment. Nature Communications, 15(1), 3936.

Smith, A. D. (1993). National Identity. University of Nevada Press.

Song, H., Park, B.-Y., Park, H., & Shim, W. M. (2021). Cognitive and Neural State Dynamics of Narrative Comprehension. The Journal of Neuroscience: The Official Journal of the Society for Neuroscience, 41(43), 8972–8990.

van Baar, J. M., & FeldmanHall, O. (2022). The polarized mind in context: Interdisciplinary approaches to the psychology of political polarization. The American Psychologist, 77(3), 394–408.

van Baar, J. M., Halpern, D. J., & FeldmanHall, O. (2021). Intolerance of uncertainty modulates brain-to-brain synchrony during politically polarized perception. Proceedings of the National Academy of Sciences of the United States of America, 118(20). 10.1073/pnas.2022491118

van Veen, V., Krug, M. K., Schooler, J. W., & Carter, C. S. (2009). Neural activity predicts attitude change in cognitive dissonance. Nature Neuroscience, 12(11), 1469–1474.

Vollberg, M. C., & Cikara, M. (2018). The neuroscience of intergroup emotion. Current Opinion in Psychology, 24, 48–52.

Waldfogel, H. B., Sheehy-Skeffington, J., Hauser, O. P., Ho, A. K., & Kteily, N. S. (2021). Ideology selectively shapes attention to inequality. Proceedings of the National Academy of Sciences of the United States of America, 118(14), e2023985118.

Westen, D., Blagov, P. S., Harenski, K., Kilts, C., & Hamann, S. (2006). Neural bases of motivated reasoning: an FMRI study of emotional constraints on partisan political judgment in the 2004 U.S. Presidential election. Journal of Cognitive Neuroscience, 18(11), 1947–1958.

Wu, Y.-S. (2005). Taiwan’s domestic politics and cross-strait relations. China Journal (Canberra, A.C.T.), 53, 35–60.

Yang, E., Milisav, F., Kopal, J., Holmes, A. J., Mitsis, G. D., Misic, B., Finn, E. S., & Bzdok, D. (2023). The default network dominates neural responses to evolving movie stories. Nature Communications, 14(1), 4197.

Yang, F.-C. I., & Mak, S. L. M. (2021). Indigenism as a project: language politics and the hegemony of postcolonialism in Taiwan. Inter-Asia Cultural Studies, 22(4), 473–493.

Yeshurun, Y., Nguyen, M., & Hasson, U. (2021). The default mode network: where the idiosyncratic self meets the shared social world. Nature Reviews. Neuroscience, 22(3), 181–192.

You, K. (2020). A study on the relationship between users’ emotional engagement in political entertainment and their political participation, using computational sentiment analysis. The Korean Society of Culture and Convergence, 42(7), 359–393.

Zebarjadi, N., Kluge, A., Glerean, E., Tassinari, M., Jääskeläinen, I. P., Jasinskaja-Lahti, I., & Levy, J. (2026). Polarized neural responses to political narratives are sensitive to small variations in self-reported political perspectives. iScience, 29(1), 114268.

Zheng, G., & Han, S. (2025). Distinct psychological and neural constructs of patriotism and nationalism. Cerebral Cortex (New York, N.Y.: 1991), 35(9). 10.1093/cercor/bhaf268

Zhong, Y. (2016). Explaining national identity shift in Taiwan. Dang Dai Zhongguo [The Journal of Contemporary China], 25(99), 336–352.

Zmigrod, L. (2021). A neurocognitive model of ideological thinking. Politics and the Life Sciences: The Journal of the Association for Politics and the Life Sciences, 40(2), 224–238.

Zmigrod, L. (2022). A psychology of ideology: Unpacking the psychological structure of ideological thinking. Perspectives on Psychological Science: A Journal of the Association for Psychological Science, 17(4), 1072–1092.

Zmigrod, L., Eisenberg, I. W., Bissett, P. G., Robbins, T. W., & Poldrack, R. A. (2021). The cognitive and perceptual correlates of ideological attitudes: a data-driven approach. Philosophical Transactions of the Royal Society of London. Series B, Biological Sciences, 376(1822), 20200424.

Zmigrod, L., Rentfrow, P. J., & Robbins, T. W. (2018). Cognitive underpinnings of nationalistic ideology in the context of Brexit. Proceedings of the National Academy of Sciences of the United States of America, 115(19), E4532–E4540.

Zmigrod, L., & Tsakiris, M. (2021). Computational and neurocognitive approaches to the political brain: key insights and future avenues for political neuroscience. Philosophical Transactions of the Royal Society of London. Series B, Biological Sciences, 376(1822),

